# Annexin A1 drives macrophage skewing towards a resolving phenotype to accelerate the regeneration of muscle injury through AMPK activation

**DOI:** 10.1101/375709

**Authors:** Simon McArthur, Thomas Gobbetti, Gaëtan Juban, Thibaut Desgeorges, Marine Theret, Julien Gondin, Juliana E Toller-Kawahisa, Christopher P Reutelingsperger, Mauro Perretti, Rémi Mounier

**Affiliations:** Institute of Dentistry, Barts & the London School of Medicine & Dentistry, Queen Mary, University of London, Blizard Institute, Newark Street, London E1 2AT, United Kingdom; William Harvey Research Institute, Barts & the London School of Medicine & Dentistry, Queen Mary, University of London, Charterhouse Square, London EC1M 6BQ, United Kingdom; Institut Neuromyogène, CNRS UMR5310, INSERM U1217, Université Claude Bernard Lyon1, France; Department of Biochemistry & Immunology, Ribeirao Preto Medical School, University of Sao Paulo, Avenida Banderiantes, 3900, Ribeirao Preto, Sao Paulo 14049-900, Brazil; Department of Biochemistry, Maastricht University, Cardiovascular Research Institute Maastricht, Maastricht, The Netherlands; Centre for Inflammation and Therapeutic Innovation, Queen Mary, University of London

**Keywords:** macrophage switch, resolution of inflammation, skeletal muscle homeostasis

## Abstract

Understanding the circuits that promote an efficient resolution of inflammation is crucial to deciphering the molecular and cellular processes required to promote tissue repair. Macrophages play a central role in the regulation of inflammation, resolution and repair/regeneration. Using a model of skeletal muscle injury and repair, herein we identify Annexin A1 (AnxA1) as the extracellular trigger of macrophage skewing towards a pro-reparative phenotype. Brought into the injured tissue initially by migrated neutrophils, and then over-expressed in infiltrating macrophages, AnxA1 activates FPR2/ALX receptors and the downstream AMPK signalling cascade leading to macrophage skewing, dampening of inflammation and regeneration of muscle fibres. Mice lacking AnxA1 in all cells or in myeloid cells only display a defect in this reparative process. *In vitro* experiments recapitulated these properties, with AMPK null macrophages lacking AnxA1-mediated polarization. Collectively, these data identify the AnxA1/FPR2/AMPK axis as a novel pathway in skeletal muscle injury regeneration.

## Introduction

An efficient inflammatory response is a necessary component of the reaction to injury or infection, but it is equally critical for this inflammatory response to be terminated in a timely and appropriate manner, enabling the restoration of tissue homeostasis (Perretti et al., 2017; Serhan, 2014). Indeed, chronic inflammation that results from failure of resolution represents a major contributing factor to a multitude of pathologies, from arthritis to sepsis to dementia (Perretti et al., 2015; Tabas and Glass, 2013).

An inflammatory reaction consists of the co-ordinated activity of numerous cells and soluble mediators, but a central role is played by macrophages. These cells, whether resident in the tissue or recruited from circulating monocyte populations, are amongst the first responders to pathogen- or damage-associated molecular patterns, initiating endothelial activation and neutrophil recruitment (Davies and Taylor, 2015). Beyond their roles as sentinels, macrophages are important drivers of the progression of an inflammatory response, acting to clear pathogens, effete cells and debris by phagocytosis (Mantovani et al., 2013), serving as antigen presenting cells to recruit the adaptive arm of the immune response (Motwani and Gilroy, 2015), and ultimately enabling processes of tissue repair and resolution (Lucas et al., 2010; Troidl et al., 2009). That one cell type is able to achieve this diverse array of functions is due in large part to their remarkable degree of phenotypic plasticity, with macrophages existing in a wide variety of forms along a spectrum running from a largely pro-inflammatory state, often indicated as M1, to a primarily non-phlogistic and pro-resolving phenotype, termed M2 (Murray et al., 2014).

A number of factors promoting this phenotypic transformation have been studied including exposure to anti-inflammatory cytokines such as interleukin (IL)-10 and IL-4, and the phagocytic removal of cell debris, although a complete description of the underlying mechanisms is lacking (Martinez and Gordon, 2014). We and others have identified a key role for the intracellular signalling pathway governed by AMP-activated protein kinase (AMPK) (Chan et al., 2015; Mounier et al., 2013; Park et al., 2017). Activation of this pathway is required for efficient conversion of pro- to anti-inflammatory-type macrophages, and inhibition of such a response significantly attenuated recovery in a model of inflammatory skeletal muscle injury (Mounier et al., 2013). Whilst AMPK is undoubtedly important in the phenotypic conversion of macrophages during the course of an inflammatory reaction, the nature of the extracellular trigger(s) for its stimulation remain unclear.

The protein annexin A1 (ANXA1) (Lim and Pervaiz, 2007; Perretti and D’Acquisto, 2009), is a major driver of inflammatory resolution, promoting neutrophil apoptosis (Solito et al., 2003), non-phlogistic monocyte recruitment (McArthur et al., 2015) and macrophage efferocytosis (Dalli et al., 2012; Scannell et al., 2007). Moreover, we and others have recently provided evidence showing ANXA1 to promote an anti-inflammatory macrophage phenotype in *in vitro* models of rheumatoid arthritis (Rhys et al., 2018) and tumour growth (Moraes et al., 2017), but how this translates to an *in vivo* situation is less clear. Against this background, we applied a well-characterised model of skeletal muscle injury (Arnold et al., 2007; Varga et al., 2013; Varga et al., 2016a) to test the hypothesis that ANXA1 and its receptor FPR2/ALX could be the upstream trigger of AMPK activation, and hence be a major driver of the pro- to anti-inflammatory macrophage phenotype shift, promoting inflammatory resolution and the restoration of skeletal muscle tissue homeostasis.

## Results

### Annexin A1 and Fpr2/3 null mice show impaired recovery from skeletal muscle injury

To investigate the role of ANXA1 and its receptor FPR2/ALX (in humans, or the orthologue Fpr2/3 in mice) in the control of macrophage phenotype we first utilised the murine model of cardiotoxin-induced *Tibialis Anterior* (TA) injury, a model characterised by necrotic tissue damage and extensive macrophage activity (d’Albis et al., 1988; Mounier et al., 2013). Analysis of immune cell infiltration in lesioned wild-type mice revealed a typical inflammatory profile, dominated by Ly6G^+^ neutrophils for the initial post-lesioning period, supplanted by F4/80^+^ macrophages from day 2 onwards (Suppl Fig 1A-B). Monitoring ANXA1 expression in the tissue revealed that while absent in uninjured tissue, the protein was primarily restricted to immune cells until day 2 post-lesioning, with regenerating muscle fibres displaying weak immunostaining from Day 7 (Suppl Fig 1A-B), as reported previously (Bizzarro et al., 2012). Whilst F4/80^+^ murine macrophages expressed AnxA1 at all time-points examined (Suppl Fig 1B-C), the proportion of macrophages expressing its primary receptor, Fpr2/3, decreased over time, from approximately 95% at day 2 to 70% at day 7 and just 5% two weeks post-lesioning, even though significant numbers of macrophages were still detected in the tissue (Suppl Fig 2A-B). Importantly, expression of Fpr2/3 on muscle fibers was not apparent at any time-point examined (Suppl Fig 2A).

Whilst uninjured TA muscle weights were comparable between wild-type, AnxA1^-/-^ and Fpr2/3^-/-^ mice (Fig 1A-B, Suppl Fig 3A), recovery of muscle mass following administration of cardiotoxin was significantly reduced in AnxA1^-/-^ and Fpr2/3^-/-^ mice than in wild-type animals (Fig 1B). Histological analysis of muscles 28 days post-injury revealed significantly reduced myofiber cross-sectional area (CSA) and myonuclei *per* fiber (*i.e*. the result of differentiation and fusion of muscle stem cells) in AnxA1^-/-^ and Fpr2/3^-/-^ animals (Fig 1C-E). These outcomes are suggestive of impaired skeletal muscle regeneration, a notion confirmed by marked lipid accumulation in AnxA1^-/-^ mice (Fig 1F). Analysis of immune cell infiltration by flow cytometry revealed that 2 days following cardiotoxin administration, approximately 60% of macrophages expressed a pro-resolving, anti-inflammatory phenotype (CD45^+^Ly6C/G^neg^F4/80^hi^) in wild-type mice, whilst AnxA1^-/-^ and Fpr2/3^-/-^ animals showed comparable levels of pro-(CD45^+^Ly6C/G^hi^F4/80^low^) and anti-inflammatory (CD45^+^Ly6C/G^low^F4/80^hi^) cells (Fig 1G, Suppl Fig 3B). Consequently, the resolution index (ratio of anti- to pro-inflammatory macrophages) of AnxA1^-/-^ and Fpr2/3^-/-^ mice was significantly lower than that of wild-type animals, indicative of a prolonged inflammatory response (Fig 1H). These results indicate that a proportion of pro-inflammatory macrophages fail to convert to an anti-inflammatory phenotype at the time of resolution in the absence of AnxA1 or Fpr2/3.

**Figure 1.**
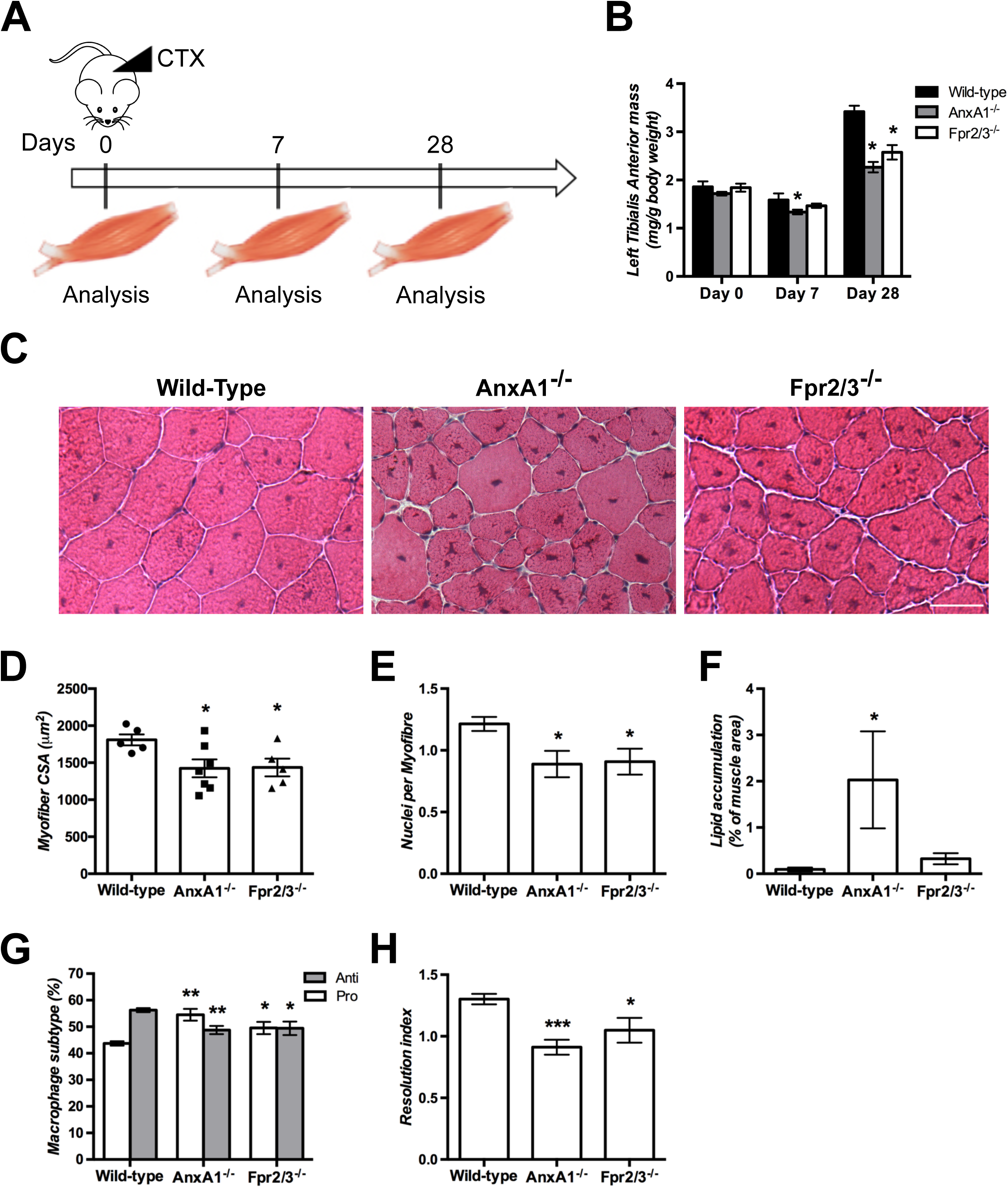
Non-redundant role of ANXA1 in cardiotoxin-induced muscle injury and repair. (A) Experimental set-up. Acute injury was induced by cardiotoxin (CTX) injection in the *Tibialis Anterior* (TA) of Wild-Type, AnxA1^-/-^ and Fpr2/3^-/-^ mice. Muscles were analysed 0, 7 and 28 days after injury. (B) TA mass normalised to mouse body weight. (C) Hematoxylin-eosin (HE) staining of muscles 28 days after injury. White bar = 50 μm. (D-E) Myofiber cross-sectional area (D), number of nuclei *per* myofiber (E) and lipid accumulation (F) in muscles 28 days post-CTX injury. (G-H) Macrophage subtypes analysis 2 days post-CTX injury. Shown are the percentage of pro- and anti-inflammatory macrophages within the F4/80^+^ population (G) and the resolution index (H). Results are mean ± SEM of at least three animals. *p<0.05, **p<0.01 and ***p<0.001 *versus* Wild-Type.

### Macrophages are the major source of AnxA1 in lesion recovery

The more pronounced effects of cardiotoxin in AnxA1^-/-^ and Fpr2/3^-/-^ mice indicate the existence of important regulatory functions for this pathway in skeletal muscle injury. However, while Fpr2/3 expression appears restricted to immune cells infiltrating the TA muscle (Suppl Fig 2), AnxA1 is expressed markedly by immune cells and to a lower extent by regenerating muscle fibres (Suppl Fig 1). As such, we deemed it important to define the primary cell source of this mediator in the repair process, a question we addressed using chimeric mice bearing wild-type muscle but AnxA1^-/-^ bone marrow-derived leukocytes. Therefore, CX3CR1-GFP mice, which harbour GFP-expressing monocytes/macrophages were irradiated and transplanted with wild-type or AnxA1^-/-^ derived bone marrow cells, prior to injection of cardiotoxin in the TA muscle (Fig 2A). Analysis of bone marrow populations at the time of euthanasia showed less than 1% of monocytes (CD115^pos^ cells) expressed GFP, suggesting >99% engraftment efficiency with either wild-type or AnxA1^-/-^ bone marrow (Fig 2B-C). Weight recovery was similar after wild-type or AnxA1^-/-^ transplant (Suppl Fig 3C), but histological analysis of TA muscles 28-days post-CTX injury revealed that animals receiving AnxA1^-/-^ bone marrow cells displayed a significantly reduced myofiber cross-sectional area than animals receiving wild-type bone marrow cells (Fig 2D-E). These data indicate that the defective regeneration quantified in AnxA1^-/-^ muscle is a consequence of an intrinsic defect in myeloid, rather than stromal, cells.

**Figure 2.**
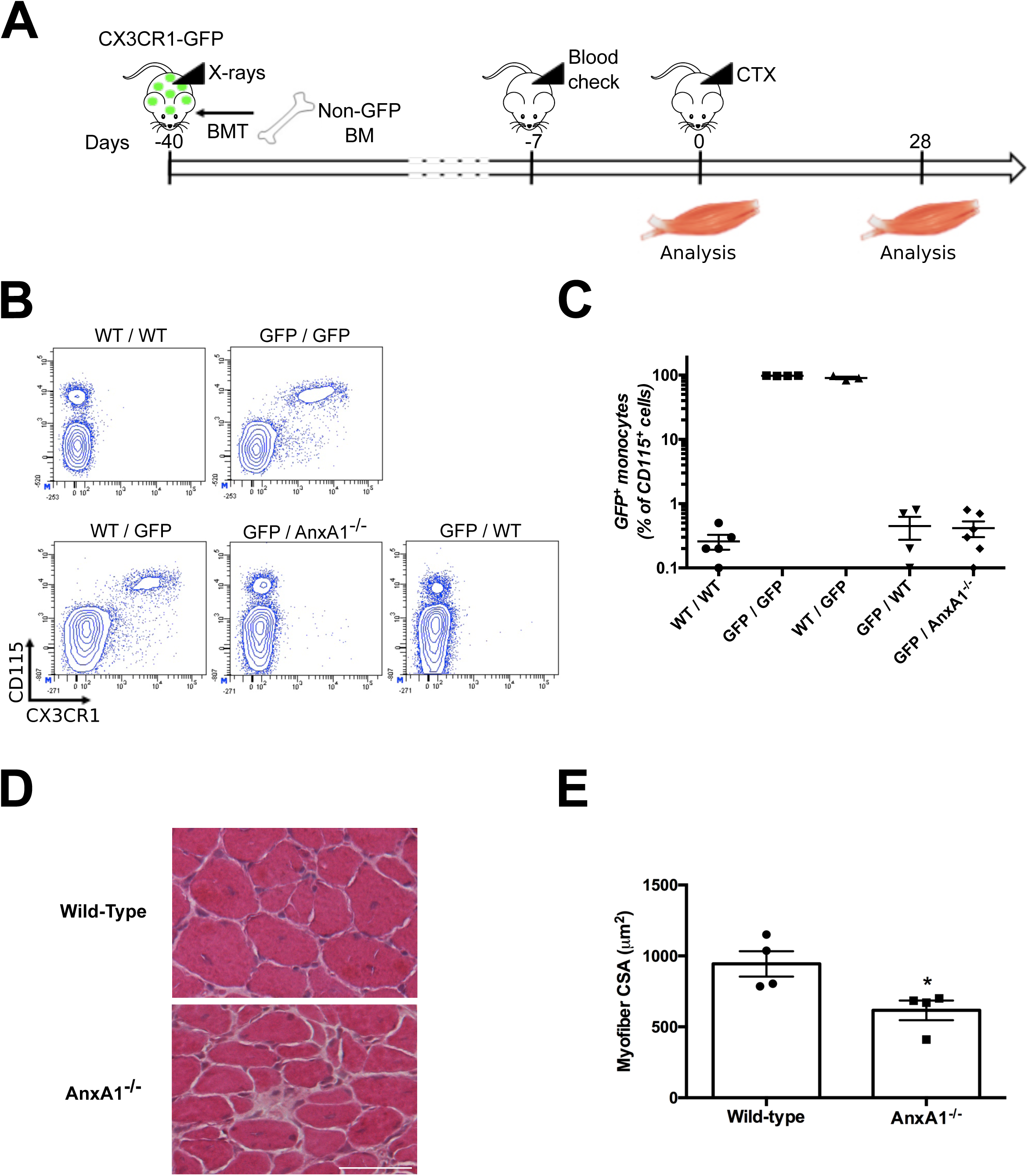
Infiltrating myeloid cell-derived ANXA1 controls muscle repair. (A) Experimental set-up. CX3CR1-GFP mice were irradiated and then transplanted with bone marrow cells isolated from Wild-Type or AnxA1^-/-^ mice. Bone marrow engraftment was checked on a blood sample after around 5 weeks. Then animals were injured in their *Tibialis Anterior* by cardiotoxin (CTX) injection and muscles analysed 0 or 28 days later. Engraftment was confirmed on the bone marrow of each animal on the day of sacrifice. (B-C) Representative FACS plots (B) and quantification (C) of the GFP^+^ monocytes in the bone marrow of the sacrificed animals. (D-E) HE staining (D) and myofiber cross-sectional area (E) of TA muscles 28 days post-CTX injury. White bar = 50 μm. Results are mean ± SEM of at least four muscles. *p<0.05 *versus* Wild-Type.

### Exogenous ANXA1 can induce human macrophage phenotype conversion in vitro

As our murine analyses indicate a potential role for AnxA1 in the polarization of macrophages from a pro-inflammatory to a pro-resolving/reparative phenotype, we investigated these effects *in vitro* using human PBMC-derived macrophages. An M1-like macrophage phenotype was induced by 24 h incubation with bacterial lipopolysaccharide (LPS) and γ-interferon (IFNγ), prior to addition of human recombinant ANXA1 (hrANXA1). Following hrANXA1 treatment, analysis of cell surface markers revealed a significant reduction in expression of the M1 marker protein major histocompatibility complex II (MHCII) (Fig 3A), accompanied by a significant increase in expression of the M2 marker protein CD206 (Fig 3B). These surface marker changes were mirrored by changes at the transcriptional level, with hrANXA1 treatment inducing a reduction in mRNA expression for the pro-inflammatory species *Tnfa* and *Nos2* paired with increased message of *Il-10* (Fig. 3C-E). No significant changes in *Tgfb1* expression were quantified (Fig. 3F). Together, these *in vitro* data support our *in vivo* findings, providing evidence that ANXA1 application favours a pro-resolving macrophage phenotype.

**Figure 3.**
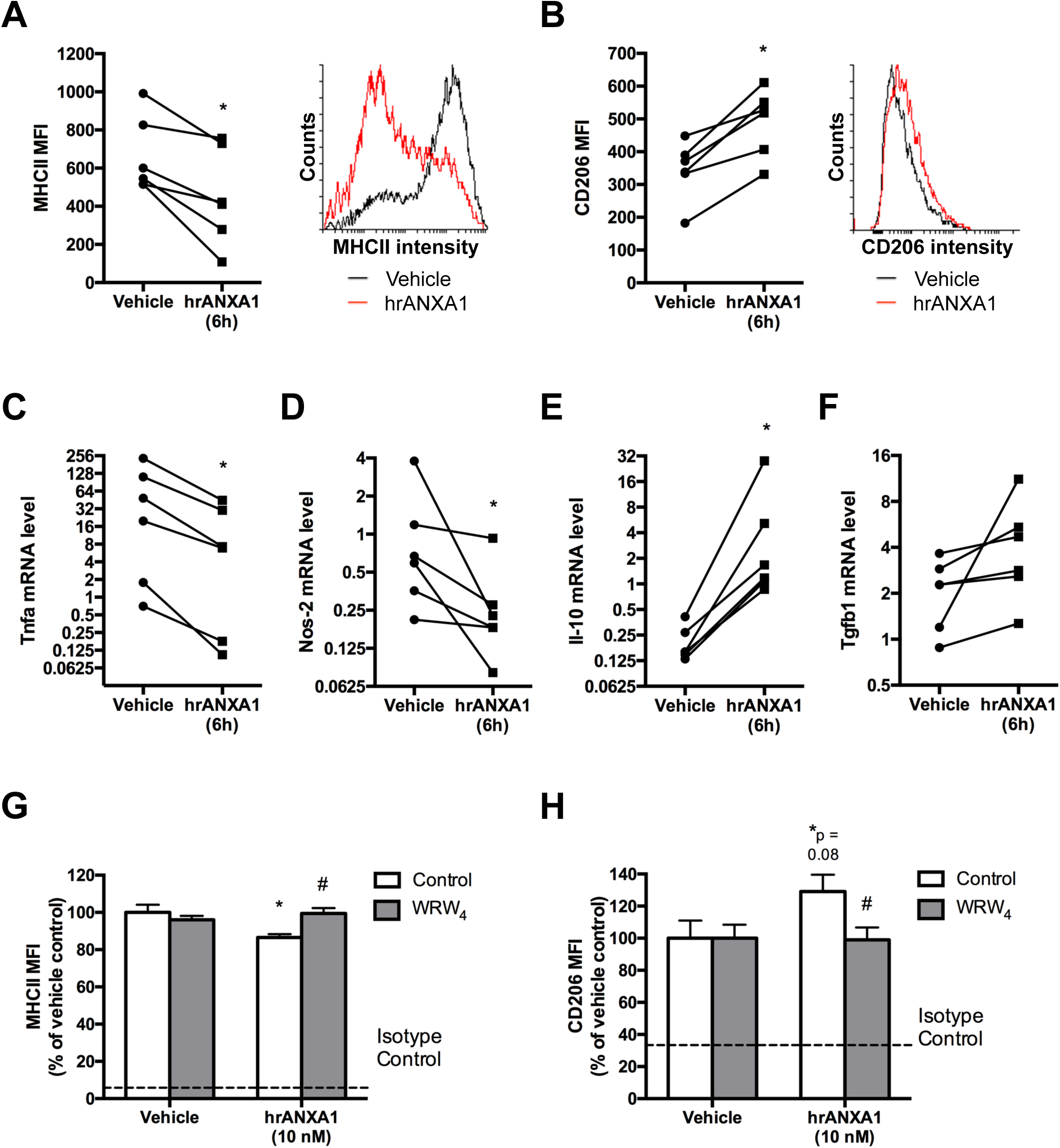
Exogenous hrANXA1 controls human and mouse macrophage polarization *in vitro*. Human PBMC-derived macrophages were incubated for 24 hours with LPS+IFNγ to promote an M1-like phenotype, prior to addition of human recombinant ANXA1 (hrANXA1, 10nM) for further 6 hours. (A-B) Mean Fluorescent Intensity (MFI) units measured by flow cytometry of MHCII pro-inflammatory (A) and CD206 anti-inflammatory (B) markers. Shown are MFI quantification (left panel) and representative FACS plots (right panel). (C-F) RT-qPCR analysis of Tnfa (C) and Nos-2 (D) pro-inflammatory genes, and Il-10 (E) and Tgfb1 (F) anti-inflammatory genes. (G-H) MFI units as measured by flow cytometry of MHCII pro-inflammatory (G) and CD206 anti-inflammatory (H) markers after treatment by hrANXA1 in the presence or absence of the FPR2/ALX antagonist WRW_4_ (10 μM). Results are mean ± SEM of at least three independent experiments. *p<0.05 *versus* Vehicle. #p<0.05 *versus* Control.

The specificity of hrANXA1 action through its human receptor FPR2/ALX was then determined, to complement the *in vivo* observations made with Fpr2/3^-/-^ mice lacking the orthologue of the human FPR2/ALX (Dufton et al., 2010). Intriguingly, we observed significant FPR2/ALX surface expression both in unstimulated human PBMC-derived macrophages and M1-phenotype cells, yet cell surface expression of the receptor was lost on cells stimulated towards an M2 phenotype with IL-4 (Suppl Fig. 4). These data are in agreement with the *in vivo* observation that Fpr2/3^-/-^ is absent from macrophages that have infiltrated the injured muscle at Day 7 and beyond, that is, when a reparative cell phenotype has been acquired (Suppl Fig 2B-C). The functional engagement of FPR2/ALX was then confirmed through use of the selective antagonist WRW_4_. The modulatory effects of hrANXA1 on pro-inflammatory PBMC-derived macrophages (*i.e.* reduced surface MHCII expression and augmented surface CD206 expression) were lost in the presence of WRW_4_ (Fig. 3G-H).

Together these experiments represent a clear *in vitro* counterpart to the effects observed in the muscle injury model, showing the key role of ANXA1 acting through FPR2/ALX to drive a shift in macrophage phenotype towards a pro-resolving and reparative polarization.

### Exogenous ANXA1 treatment stimulates AMPK activation through FPR2/ALX

The enzyme 5’-adenosine monophosphate-activated protein kinase, AMPK, plays a critical role in macrophage phenotype skewing and is necessary for efficient regeneration after skeletal muscle damage (Mounier et al., 2013). We queried whether this signalling pathway would underlie the effects of hrANXA1 upon human macrophage phenotype and indeed muscle repair *in vivo*.

Exposure of human PBMC-derived macrophages to hrANXA1 activated a number of components of the AMPK signalling pathway, promoting phosphorylation of Ca^2+^/calmodulin-dependent protein kinase II, AMPKα1 itself and its downstream effector acetyl-CoA carboxylase (Fig 4A-B). Interestingly, the phosphorylation of these proteins in macrophages only occurred to an appreciable degree following exposure to hrANXA1 for 30-60 min (Fig 4B-D), in contrast to early MAP kinase signalling previously reported in human monocytes (McArthur et al., 2015). That activation of AMPKα1 depended upon binding of hrANXA1 to FPR2/ALX was confirmed through analysis of the effects of the antagonist WRW_4_, which abrogated hrANXA1-evoked AMPKα1 phosphorylation in human macrophages (Fig 4E). Analyses of murine bone marrow-derived macrophages from wild-type and Fpr2/3^-/-^ animals confirmed the pivotal role of this receptor: whilst treatment with hrANXA1 induced phosphorylation of AMPKα1 in wild-type cells, this response was absent in macrophages from Fpr2/3^-/-^ mice (Fig 4F).

**Figure 4.**
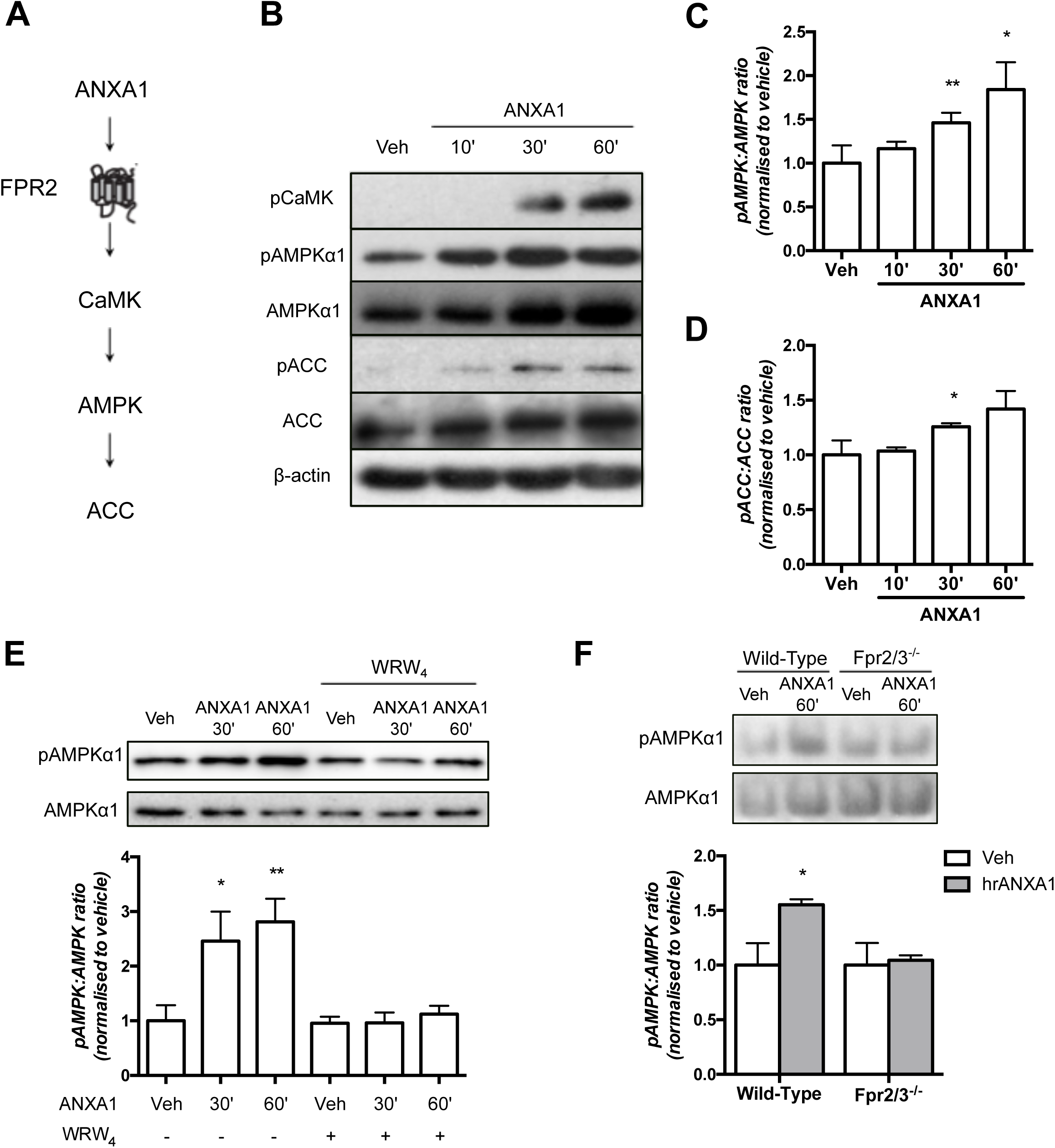
The ANXA1/FPR2/ALX pathway activates the AMPK signalling cascade in human and murine macrophages. (A) Schematic representation of the ANXA1/FPR2/ALX signalling cascade. (B-D) Western blot analysis of pCaMK, pAMPKα1 and pACC in human PBMC-derived macrophages treated with hrANXA1. Shown are representative blots (B) and quantification of pAMPKα1 to AMPKα1 (C) and pACC to ACC (D) ratios. (E) Representative western blot (top panel) and quantification (bottom panel) of pAMPKα1 to AMPKα1 ratio in human PBMC-derived macrophages treated by hrANXA1 in the presence or absence of 10 μM WRW_4_. (F) Representative western blot (top panel) and quantification (bottom panel) of pAMPKα1 to AMPKα1 ratio in Wild-Type or Fpr2/3^-/-^ murine bone marrow derived macrophages treated with hrANXA1. Results are means ± SEM of at least three independent experiments. *p<0.05, **p<0.01 *versus* Vehicle.

### AMPK activation is required for macrophage phenotype conversion induced by ANXA1

We investigated the relationship between FPR2/ALX-mediated AMPK activation and the shift in macrophage phenotype induced by hrANXA1 treatment through analyses in primary bone marrow-derived macrophages taken from wild-type mice and animals lacking the key catalytic α_1_ subunit of AMPK (Jorgensen et al., 2005). Whilst treatment of wild-type macrophages with hrANXA1 (10nM, 24h) reduced the percentage of macrophages positive for the pro-inflammatory markers iNOS and CCL3, this response was notably absent in cells from AMPKα1^-/-^ mice (Fig 5A). Correspondingly, hrANXA1 treatment augmented the proportion of wild type cells expressing the anti-inflammatory markers TGFβ1 (notably at variance from human macrophages), CD163 and CD206, but this did not occur in cells lacking AMPKα1 (Fig 5B). Together, these data indicate activation of AMPKα1 as a key step in the macrophage phenotype shift induced by ANXA1.

**Figure 5.**
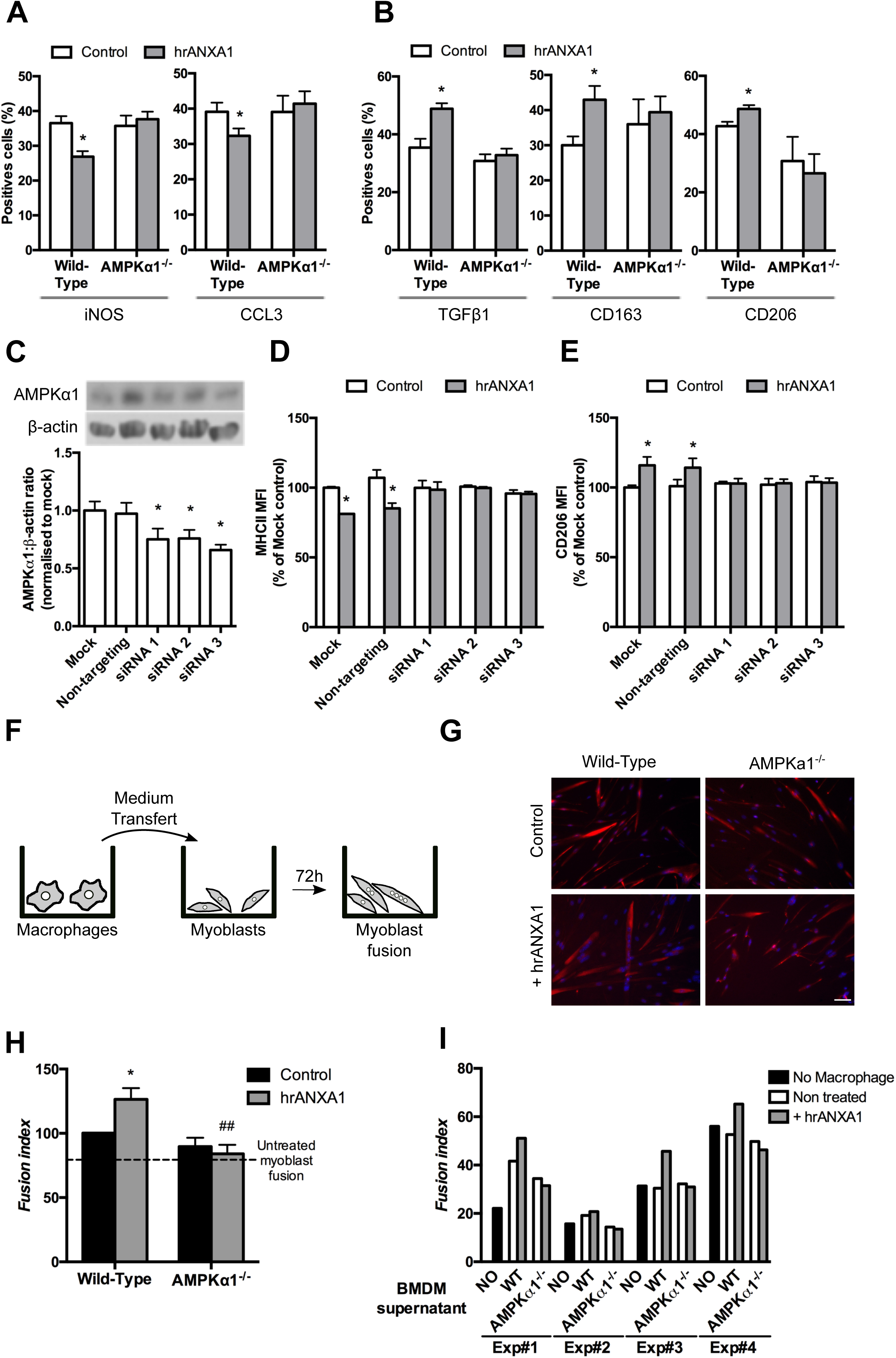
Null or reduced AMPK expression affects ANXA1-mediated macrophage polarization. (A-B) Primary macrophages derived from Wild-Type or AMPKα1^-/-^ mice were treated with 10nM hrANXA1 and the percentage of cells expressing the pro-inflammatory markers iNOS and CCL3 (A) and the anti-inflammatory markers TGFB1, CD163 and CD206 (B) was determined by immunofluorescence. (C-E) Human PBMC-derived macrophages were transfected by a non-targeting or three different AMPKα1-targeting siRNAs and treated with 10 nM hrANXA1 for 24 h. AMPKα1 protein level was determined by western blot (C) and the MFI units of the pro-inflammatory MHCII (D) and anti-inflammatory CD206 (E) markers were measured by flow cytometry. (F-I) Conditioned medium produced by murine macrophages was transferred onto murine primary myoblasts and their fusion was measured by immunofluorescence. (F) Experimental set-up. (G) Representative images of desmin (red) and Hoechst (blue) labelling of myoblast cultures. White bar = 50 μm. (H-I) Fusion index calculated after desmin labelling. Shown are the means +/- SEM of the independent experiments (H) and the results of each individual replicate (I). Results are mean ± SEM of at least three independent experiments. *p<0.05 versus Mock or Control. ^##^p<0.01 versus Wild-Type.

To confirm these findings in human cells, we employed an RNA interference approach, transfecting primary PBMC-derived macrophages with three distinct siRNA constructs targeting the AMPKα1 subunit (Fig. 5C). Treatment of M1-like pro-inflammatory macrophages for 6 h with 10 nM ANXA1 reduced expression of MHCII and augmented CD206 in mock transfected cells and in cells bearing a non-targeting siRNA construct; this effect was absent in cells transfected with any of three different siRNA constructs targeting human AMPKα1 (Fig. 5D-E).

### Adult myogenesis is ANXA1-AMPK signalling dependent in vitro

Together, these data from human and mouse macrophages make a compelling case that macrophage phenotype shifting can be induced through an ANXA1/FPR2/AMPK cascade. To investigate whether this process underlies differences in recovery from cardiotoxin-induced muscle lesions seen between wild-type and AnxA1^-/-^ and Fpr2/3^-/-^ mice, we made use of an established model of *in vitro* muscle repair (Mounier et al., 2013; Varga et al., 2016b).

We employed an *in vitro* model of adult myogenesis in which conditioned medium from primary bone marrow-derived macrophages was used to stimulate primary murine myoblasts for 72h, quantifying the proportion of multinucleated myotubes (Fig. 5F). As such this *in vitro* setting recapitulates the processes of adult myogenesis (activation, differentiation, migration and fusion of muscle cells) that occur during skeletal muscle regeneration (Mounier et al., 2013; Varga et al., 2016b). Conditioned medium from ANXA1-treated wild-type macrophages induced an increase in myotube fusion index (Fig 5G-I). This response was absent when myotubes were treated with conditioned medium from ANXA1-treated AMPKα1^-/-^ macrophages (Fig 5G-I).

## Discussion

The timely resolution of inflammation is a fundamental requirement for restoring homeostasis following infection or damage, with its failure being a significant contributory factor to numerous chronic inflammatory pathologies. Macrophages are key players in this process, given their ability to transition from generally pro-inflammatory to anti-inflammatory phenotypes (Motwani and Gilroy, 2015). Substantial effort has gone into deciphering the complex signals underpinning macrophage plasticity, with numerous soluble mediators having been implicated (Martinez and Gordon, 2014), but the mechanistic link between these factors and changes in phenotype remains poorly understood especially when investigated in specific tissue-restricted settings. In the current study, we have employed a well-characterised model of muscular injury and recovery (Mounier et al., 2013) to identify the ability of myeloid cell-derived ANXA1 to promote an anti-inflammatory macrophage phenotype, promoting resolution and tissue repair. Moreover, we show the actions of this protein to be mediated through the cell surface receptor FPR2/ALX and consequent activation of the intracellular signalling molecule AMPK, mechanistically linking external and intracellular pro-resolving signals governing macrophage phenotype.

These data reinforce the status of the ANXA1-FPR2/ALX pair as a major endogenous driver of inflammatory resolution, and add to its known roles in regulating neutrophil apoptosis (Solito et al., 2003), efferocytosis (Maderna et al., 2005; Scannell et al., 2007), and the recruitment of monocytes to inflammatory sites (Gobbetti et al., 2014; McArthur et al., 2015). Notably, despite muscle tissue itself beginning to express ANXA1 during tissue repair, as noted previously (Bizzarro et al., 2012), experiments with chimeric mice confirmed that the most important source of the protein to enable muscle repair remains the myeloid cells themselves. While the current studies do not permit conclusive determination of whether neutrophils or macrophages are the primary source of endogenous ANXA1 in the injured tissue, neutrophils contain substantial quantities of ANXA1 (Francis et al., 1992), which we have previously shown to be a major monocyte chemoattractant (McArthur et al., 2015) suggesting that the protein - produced at an early stage by and released from infiltrating neutrophils (Damazo et al., 2006) - may act to both recruit monocytes and promote a pro-resolving phenotype in resulting macrophages. Whatever the ultimate cellular source of ANXA1 however, the dependency upon the presence of ANXA1-expressing leukocytes for efficient macrophage phenotypic conversion is another example of how “the beginning programs the end” in resolution (Serhan and Savill, 2005), emphasising the finely tuned nature of the acute inflammatory response and its ability to encode its own termination.

Besides highlighting the role of ANXA1 as a regulator of inflammatory resolution (Leoni and Nusrat, 2016), these new data add further weight to the importance of its principal receptor FPR2/ALX in this process, identifying it as a conduit for induction of a pro-resolving macrophage phenotype through AMPK activation. It is notable that macrophage FPR2/ALX expression declined during the course of the response to muscular injury in our study; similarly, FPR2/ALX expression was significantly lower on M2 when compared with M1 phenotype cells *in vitro*. This change in receptor expression may reflect a strategy whereby once the macrophage is polarised towards a reparative phenotype, the utility of the FPR2/ALX signalling is then limited or unnecessary.

There is considerable redundancy in the mediators known to induce the conversion of macrophage phenotypes, as is perhaps to be expected given the importance of this process to the restoration of homeostasis after infection or damage, including immune complexes, apoptotic cells and a number of cytokines (Amici et al., 2017). Notably however, a significant proportion of the soluble pro-resolving mediators so far identified derive from the adaptive arm of the immune response, particularly from T_H_2 lymphocytes (Martinez and Gordon, 2014). Our data however, highlight the ability of signalling components derived from the innate side of the immune system to promote resolution in the absence of significant activation of adaptive mechanisms. Moreover, these results support the concept of how it is crucial to decipher which mediator and signal(s) are operative in specific tissues and organs to control macrophage polarization and the overall process of resolution and repair.

Our data are the first to associate activation of FPR2/ALX and the major regulator of cellular metabolism, AMPK, showing the central involvement of FPR2/ALX-stimulated AMPK activation in the induction of a pro-resolving macrophage phenotype. The precise mechanism linking AMPK activation with a change in phenotype is as yet unclear, but there is increasing evidence that changes in the metabolic status of immune cells can affect their inflammatory activity (O’Neill et al., 2016; Pearce and Pearce, 2013). Pro-inflammatory dendritic cells and T lymphocytes are characterised by high levels of glycolysis, akin to the Warburg shift described in cancer (Rodriguez-Prados et al., 2010), whereas immune cells with an anti-inflammatory or pro-resolving profile tend to exhibit greater mitochondrial respiration (Jha et al., 2015; Mounier et al., 2013; Tannahill et al., 2013). The mechanistic details of how such changes in metabolic phenotype relate to immune function, and indeed whether these differences reflect or drive immunophenotype is unclear (Van den Bossche et al., 2017), but it is notable that AMPK is a significant promoter of mitochondrial respiration and a regulator of glycolysis through the modulation of lactate dehydrogenase activity (Theret et al., 2017), driven by its ability to respond to energy debt and an increased AMP:ATP ratio (Mounier et al., 2015). Activation of this pathway would therefore seem ideally placed to induce the metabolic phenotype most closely associated with anti-inflammatory macrophage activation, but this hypothesis requires further investigation.

AMPK is involved in different cellular mechanisms that regulate skeletal muscle homeostasis (Kjobsted et al., 2018). Our previous findings highlight the importance of crosstalk between AMPK and the mTOR signaling pathway for the control of muscle cell size in the adaptive response of skeletal muscle (Lantier et al., 2014; Lantier et al., 2010; Mounier et al., 2011; Mounier et al., 2009). Moreover, we have recently shown that AMPKα1, activated following phagocytosis, is crucial for macrophage skewing from a pro- to an anti-inflammatory phenotype during resolution (Mounier et al., 2013), demonstrating that the CAMKKII/AMPKα1 pathway within macrophages is required for proper and complete skeletal muscle regeneration. Anti-inflammatory macrophages promote myogenic differentiation and fusion (Saclier et al., 2013a; Saclier et al., 2013b; Varga et al., 2016b), a finding of importance for skeletal muscle muscle regeneration where a sequential presence of pro-then anti-inflammatory cells is necessary for efficient regeneration process (Arnold et al., 2007; Varga et al., 2013). Therefore, the pro-resolving effect of ANXA1-AMPK signalling in macrophages is likely to be beneficial for skeletal muscle regeneration, as suggested by the positive effect of conditioned medium from ANXA1-treated macrophages on adult myogenesis *in vitro*.

In summary, we present here a novel mechanism governing the conversion of pro-inflammatory macrophages to a pro-resolving phenotype, linking leukocyte-derived ANXA1, FPR2/ALX and intracellular AMPK activation. This finding reinforces the position of the ANXA1-FPR2/ALX pathway as a pivotal regulator of inflammatory resolution, and suggests that it may represent a suitable target for therapeutic exploitation for the innovative treatment of pathologies characterised by chronic, non-resolving inflammation.

## Acknowledgments

This work was supported by CNRS, French Society of Myology and Wellcome Trust Programme Grant 086867/Z/08/Z. GJ was supported by Fondation pour la Recherche Medicale (Equipe FRM DEQ20140329495.

## Author Contributions

SM, TG, GJ, JG, MT & RM performed TA lesion and analysis, GJ & RM produced and analyzed chimeric mice, SM, TG and JETK performed analysis of human macrophages, GJ, JG, MT & RM performed analysis of AMPKα1 null mice, TD & RM performed murine macrophages and myoblast fusion analysis, CPR produced and provided human recombinant ANXA1, SM, MP & RM conceived and designed the study, all authors contributed to the drafting of the paper.

## Declaration of Interests

## Methods

### Animals

All procedures were performed under the UK Animals (Scientific Procedures) Act, 1986 in the UK or in compliance with European legislation in France. Animal facilities are fully licensed by relevant national authorities and protocols have been validated by ethical committee. Male C57Bl/6 mice, male *alx/fpr2/3*^*GFP/GFP*^ mice (referred to as Fpr2/3^-/-^) bearing a knocked-in green fluorescent protein (Dufton et al., 2010) and male *anxA1*^-/-^ mice (Hannon et al., 2003), aged 10 weeks were used for *in vivo* experiments. Both transgenic strains were fully backcrossed onto a C57Bl/6 genetic background.

### Production of Recombinant Human Annexin A1

Human recombinant annexin A1 (hrANXA1) was produced by a prokaryotic expression system and purified essentially as described previously (Kusters *et al*, 2015). Briefly, cDNA for human ANXA1 was inserted into the expression vector pQE30Xa (Qiagen) and transfected into E.coli (#SG13009 pREP4, Novagen) which were then grown in Luria-Bertani broth medium supplemented with ampicillin (50 μg/ml, Roche), kanamycin (30 μg/ml, Thermofisher) and 0.5% glycerol. Protein overexpression was initiated by addition of 5mM isopropyl β-D-1-thiogalactopyranoside (Eurogentec) and proteins were purified by IMAC. Purity and homogeneity were assessed by SDS-PAGE, western blotting and MALDI-TOF/TOF analysis. Endotoxin was determined with the Endosafe-PTS (FDA-licensed LAL cartridge from Charles-River) according to the manufacturer’s protocol. HrANXA1 contained < 0.2 unit endotoxin *per* mg hrANXA1 protein.

### Skeletal Muscle Injury

Skeletal muscle injury was caused by intramuscular injection of cardiotoxin (Latoxan) in the *Tibialis Anterior* (TA) muscle of male animals, as described previously (Mounier et al., 2013). Left TA muscles were injected with cardiotoxin (50 μl *per* TA, 12 μM); 1, 2, 7, 14 or 28 days post-lesioning animals were killed by exposure to CO_2_. TA were isolated and snap-frozen in liquid nitrogen-cooled isopentane for storage and later analysis.

### Murine Bone Marrow-Derived Macrophages

Bone marrow-derived macrophages (BMDMs) were prepared from adult male wild-type sv129/C57Bl6 and *Prkaa1*^-/-^ mice (referred to as AMPKα1^-/-^ (Jorgensen et al., 2005)). Mice were killed by cervical dislocation under isofluorane anaesthesia, and marrow was flushed from tibiae and femurs. Cells were plated, washed and grown for 6-7 days in DMEM High Glucose High Pyruvate, 20% heat inactivated fetal calf serum (ThermoFisher), 30% L929-cell conditioned medium, 1% Amphotericin B (2.5 μg/ml, ThermoFisher) and 100 μg/ml streptomycin (ThermoFisher). For phenotypic characterisation, BMDMs were fixed for 10 minutes in 4% formaldehyde, permeabilized for 10 minutes in PBS with 0.5 % Triton X-100 and blocked for 1 hour in PBS with 4 % BSA. They were then labeled overnight at 4°C with anti-NOS2 (#ab15323, Abcam), anti-CCL3 (#ab32609, Abcam), anti-TGFβ1 (#ab64715, Abcam), anti-CD163 (#sc-20066, Santa-Cruz) and anti-CD206 (#sc-58987, Santa-Cruz), followed by incubation for 1 hour at 37°C with FITC- or Cy3-conjugated secondary antibodies (Jackson Immunoresearch Inc). Cells were stained with Hoechst (Sigma) and mounted in Fluoromount (Interchim) and pictures were taken on an Axio Imager.Z1 (Zeiss) at 20X of magnification connected to a CoolSNAP MYO CCD Camera (Photometrics).

### *In vitro* model of adult myogenesis

Macrophages were obtained from bone marrow (BM) precursor cells that were cultured in DMEM containing 20% FBS and 30% conditioned medium of L929 cell line (enriched in CSF-1) for 7 days. Macrophages were activated with human recombinant annexin A1 for 3 days (10 nM) in DMEM containing 10% FBS. After the washing steps, serum-free DMEM was added for 24 hr to obtain macrophage-conditioned medium. Murine myogenic precursor cells (MPCs) were obtained from TA muscle and cultured using standard conditions in DMEM/F12 medium (Gibco Life Technologies) containing 20% heat inactivated Foetal Bovine Serum (FBS) and 2% G/Ultroser (Pall Inc). MPCs were seeded at 30,000 cell/cm^2^ on Matrigel (diluted 1:10) and incubated for 3 days with conditioned medium containing 2% heat inactivated horse serum. Cells were then incubated with an anti-desmin antibody (#ab32362, Abcam), followed by a Cy3-conjugated secondary antibody (Jackson Immunoresearch Inc) (Mounier *et al*, 2013; Varga *et al*, 2016). Cells were stained with Hoechst (Sigma) and mounted in Fluoromount (Interchim) and pictures were taken on Axio Observer.Z1 (Zeiss) at 20X of magnification connected to a CoolSNAP HQ2 CCD Camera (Photometrics

### Bone Marrow Transplantation

Total bone marrow cells were isolated by flushing the tibiae and femurs of 8- to 20-week-old donor mice (wild-type or AnxA1^-/-^ males) with RPMI-1640 / 10% FBS. They were transplanted into 8- to 12- week-old recipient CX3CR1-GFP^+/-^ males (monocytes/macrophages expressing GFP) previously lethally irradiated for 10 min with a dose of 0.85 Gy/min in an X-RAD 320 (Precision X-Ray). Total bone marrow cells were injected (10^7^ cells diluted in 100 μL of RPMI-1640 / 50 % mouse serum) into the retro-orbital vein of recipient mice. After transplantation, mice were fed with ciprofloxacin (10 mg/kg/day) in the drinking water for 3-weeks. Engraftment efficiency was determined on peripheral blood 5 weeks after the transplantation and on bone marrow when mice were sacrificed by FACS analysis. Briefly, red cells were lysed with ACK buffer and leukocytes were incubated with FcR Blocking Reagent (Miltenyi Biotec) for 20 min at 4°C. Finally, cells were labeled with an APC-conjugated anti-CD115 antibody for 30 min at 4°C and analysed on a BD FACS Canto II (BD Biosciences). DAPI was used as viability marker. Engraftment was determined as the percentage of monocytes not expressing GFP.

### Human Peripheral Blood-Derived Macrophages

Human cells were prepared according to an approved protocol (East London & the City Local Research Ethics Committee; no. 06/Q605/40; P/00/029). Peripheral blood was collected from healthy volunteers by intravenous withdrawal in 3.2% sodium citrate solution (1:10). Peripheral blood mononuclear cells were isolated by density centrifugation on a Histopaque-1077 gradient (Sigma) according to the manufacturer’s instructions, and were plated in RPMI 1640 for 1 hour. Cells were washed three times with ice-cold PBS without Ca^2+^/Mg^2+^ to remove lymphocytes, and adherent cells were incubated in RPMI 1640 containing 20% heat inactivated fetal calf serum for 14 days.

### Histological and Immunohistochemical Analysis

For histological analysis, muscles were harvested, snap frozen in liquid nitrogen-chilled isopentane and kept at -80°C until use. Cryosections (10μm) were prepared for hematoxylin-eosin (HE) or Sudan Black staining. Fluorescence immunohistochemical analysis was performed according to standard procedures. Briefly, transverse muscle cryosections (10 μm) were post-fixed by incubation for 15 minutes in 4% formaldehyde, blocked and immunostained using primary antibodies directed against Ly6G (1:100; #127602, Biolegend), F4/80 (1:200, #123102, Biolegend) ANXA1 (1:1,000; #71-3400, ThermoFisher) or Fpr2/3 (1:100, #sc-18191-R, SantaCruz). Secondary antibodies were Alexa Fluor 488- or 594-conjugated goat anti-rabbit or anti-rat IgG (1:300; Invitrogen). Sections were counterstained with DAPI, mounted and examined using a TCS SP5 confocal laser scanning microscope (Leica Microsystems) fitted with 405 nm, 488 nm and 594 nm lasers, and attached to a Leica DMI6000CS inverted microscope fitted with a 40× objective lens (NA 0.75 mm; working distance, 0.66 mm). Images were captured with Leica LAS AF 2.6.1 software and analysed using ImageJ 1.51w software (National Institutes of Health).

### *In vivo* Macrophage Phenotype Analysis

Macrophage phenotype was analyzed as previously described (Mounier et al., 2013). Briefly, CD45^+^ cells were isolated from regenerating muscle TA using magnetic beads conjugated to anti-CD45 antibody (Milteny Biotec) and then incubated with Fc-block (Milteny Biotec) for 30 min at 4°C. Finally, CD45^+^ cells were stained with antibody against Ly-6C/G (#17-5931-82, eBioscience) and against F4/80 (#12-4801-82, eBioscience). Percentages of Ly-6C/G^hi^F4/80^low^ and Ly-6C/G^neg^F4/80^hi^ cells were calculated among total F4/80^pos^ cells following analysis by flow cytometry with a FACSCalibur instrument (Becton Dickinson, UK) and FlowJo v.9.2 analysis software as described below.

### AMPKα1 siRNA

Primary human PBMC-derived macrophages were transfected with one of three different commercial siRNA sequences designed to target AMPKα1 or an Allstars negative control siRNA sequence using Hiperfect transfection reagent according to the manufacturer’s instructions (final concentration 2 nM; all Qiagen GmbH, Hilden, Germany), alongside mock transfected cells. After 48 hours, cells were analysed for phenotypic conversion following hrANXA1 treatment (6 hours, 10 nM). A proportion of cells were analysed for AMPKα1 expression by western blot using a rabbit polyclonal antibody raised against human the AMPKα1 subunit (1:1,000, #2795, Cell Signalling Technology) and for β-actin expression using a mouse monoclonal antibody (1:10,000; #A5316, Sigma).

### Human Macrophage Flow Cytometry Analysis

Primary human PBMC-derived macrophages were labelled with FITC-conjugated mouse monoclonal anti-MHCII (#11-9956-42, Thermofisher) and APC-conjugated mouse monoclonal anti-CD206 (#17-2069-42, Thermofisher) or isotype controls (all Thermofisher) according to manufacturer’s protocols. In all cases, 20,000 events were acquired using a FACSCalibur flow cytometer (Becton Dickinson), and analysed using FlowJo analysis software (Version 9.2, Treestar Inc). In some cases, macrophages were analysed for surface expression of FPR2/ALX; surface FcγR were blocked by incubation for 20 minutes at 4°C with IgG block (ThermoFisher Scientific, UK), followed by incubation for 30 minutes at 4°C with mouse monoclonal anti-FPR2/ALX (1μg/106 cells; GM1D6, Aldevron, Freiburg, Germany) then incubation for 30 minutes at 4°C with secondary antibody (AF488-conjugated goat anti-mouse 1:300; ThermoFisher Scientific, UK).

### Western Blot Analysis

Samples boiled in 6× Laemmli buffer were subjected to standard SDS-PAGE (10%) and electrophoretically blotted onto Immobilon-P polyvinylidene difluoride membranes (Millipore, Watford, UK). Membranes were incubated with antibodies raised against human phospho-Ca^2+/^calmodulin-dependent kinase (#12716, Cell Signalling Technology), phospho-AMPKα1 (#2531, Cell Signalling Technology), AMPKα1 (#2795, Cell Signalling Technology), phospho-acetyl-CoA carboxylase (#3661, Cell Signalling Technology), acetyl-CoA carboxylase (#3662, Cell Signalling Technology, all 1:1000) or β-actin (1:10,000; #A5316, Sigma) in Tris-buffer saline solution containing 0.1 % Tween-20 and 5 % (w/v) non-fat dry milk overnight at 4 °C. Membranes were washed for 30 minutes with Tris-buffer saline solution containing 0.1 % Tween-20, with the solution being changed at 10 minute intervals; membranes were then incubated with secondary antibody (horseradish peroxidase–conjugated goat anti-mouse 1:5000; ThermoFisher), for 2 hours at room temperature. Proteins were then detected using the enhanced chemiluminescence detection kit and visualized on Hyperfilm (Amersham Biosciences). Films were digitised and analysed using ImageJ 1.51w software (National Institutes of Health).

### Quantitative RT-PCR

Total RNA was prepared from primary human PBMC-derived macrophages using TRIzol reagent (Life Technologies Ltd), and then reverse transcribed with superscript III reverse transcriptase (Life Technologies Ltd) according to the manufacturer’s protocols. Resultant cDNA was then analysed by real-time PCR in duplicate, using the Quantitect primer system (Primer sets: FPR2/ALX QT00204295, IL-10 QT00041685, NOS-2 QT00068740, TNF-α QT00029162 and TGFβ1 QT00000728; all Qiagen Ltd.) and Power SYBR Green PCR Master Mix (Applied Biosystems). Reactions were performed in 384 well-format using the ABI Prism 7900HT Sequence Detection System. The PCR conditions consisted of 95 °C 15 min, [95 °C 15 s - 55 °C 30 s - 72 °C 30 s] × 40, with a dissociation step [95 °C 15 s/60 °C 15 s/95 °C 15 s] included after the PCR reaction to confirm the absence of non-specific products. Data was acquired and analyzed with SDS 2.3 (Applied Biosystems); fold change was calculated as 2^-ΔΔCt^.

### Statistical Analysis

All quantified *in vitro* data are derived from at least three independent donors, with experiments performed in triplicate, and are expressed as mean ± SEM. Murine *in vivo* experiments were performed with a group size of n=6, sufficient to identify a 20% effect size with a power of 0.8, and are expressed as mean ± SEM. Data were analyzed by one- or two-way ANOVA as appropriate, with *post hoc* comparison using Tukey’s HSD test. For murine *in vitro* experiments, at least 3 independent experiments were performed and statistical significance was determined using Student’s t test. In all cases, a *P*<0.05 was taken as indicating statistical significance.

---

**Suppl Fig 1.**
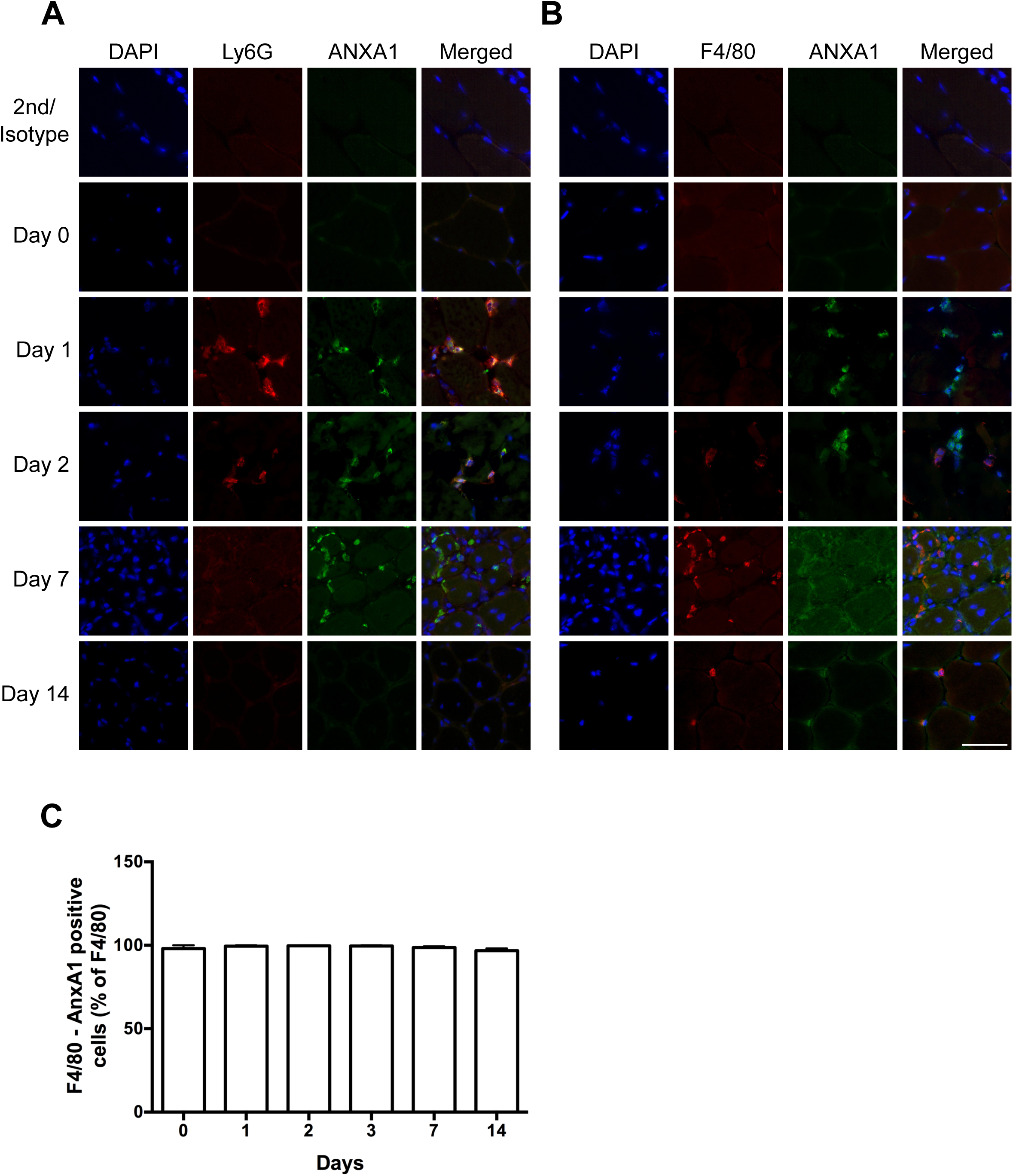
Related to Figure 1. Immune cell recruitment and pattern of AnxA1 expression in cardiotoxin-induced muscle injury and repair. Immunofluorescence analysis of ANXA1 protein in *Tibialis Anterior* muscle after cardiotoxin injury. Images show co-localisation of ANXA1 protein with Ly6G^+^ (A) and F4/80^+^ (B) cells. White bar = 50 μm. (C) Quantification of F4/80^+^ ANXA1^+^ cells in *Tibialis Anterior* (TA) muscle after cardiotoxin (CTX) injury.

**Suppl Fig 2.**
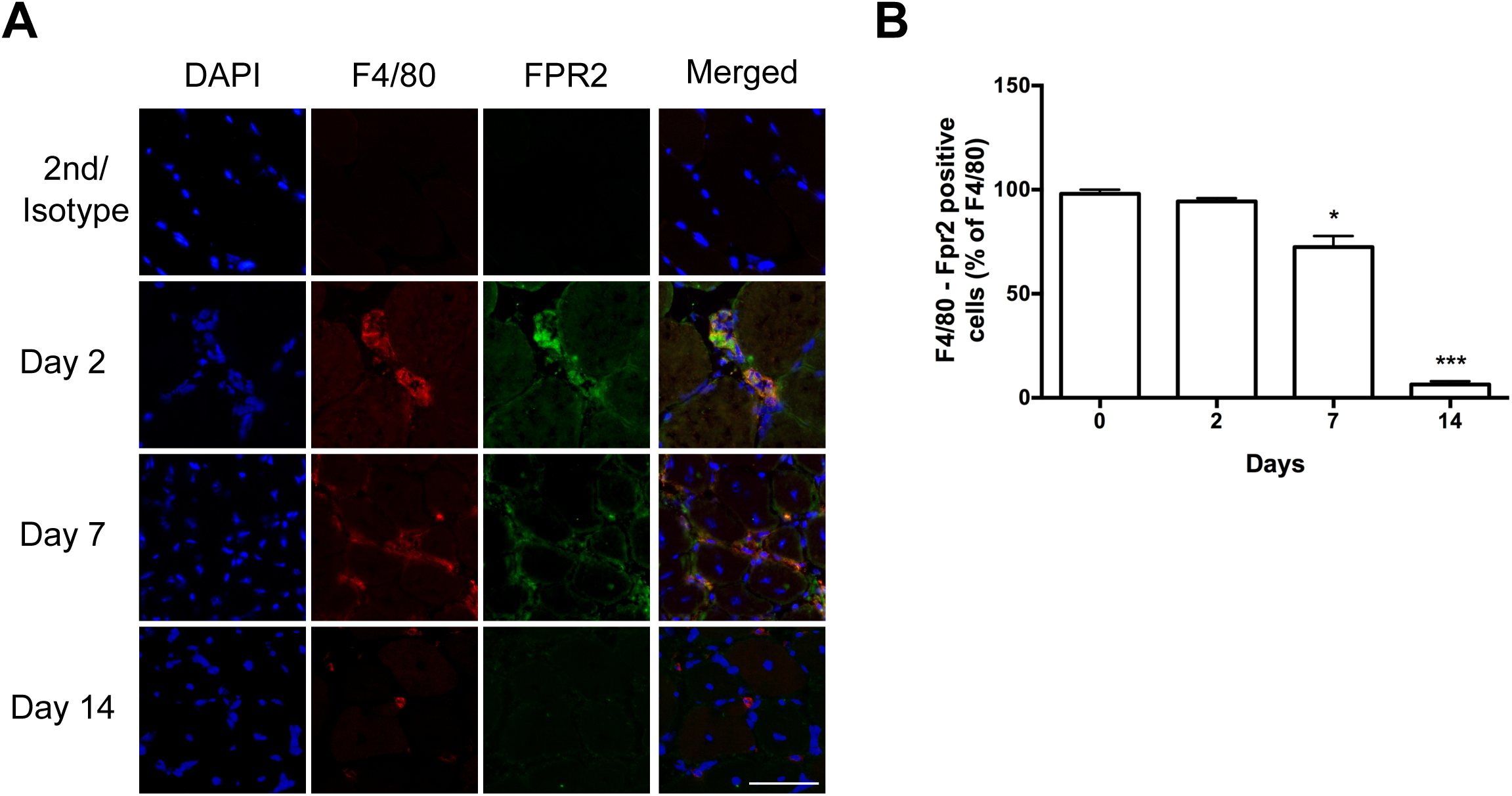
Related to Figure 1. Immune cell recruitment and Fpr2/3 expression in cardiotoxin-induced muscle injury and repair. Quantification of F4/80^+^ ANXA1^+^ cells in *Tibialis Anterior* (TA) muscle after cardiotoxin (CTX) injury (A), representative immunofluorescence analysis of Fpr2/3 protein in TA muscle after cardiotoxin (CTX) injury (B) and quantification of the percentage of F4/80^+^ cells expressing Fpr2/3 (C). White bar = 50 μm. Results are means ± SEM of three animals. *p<0.05 and ***p<0.001 *versus* Day 0.

**Suppl Fig 3.**
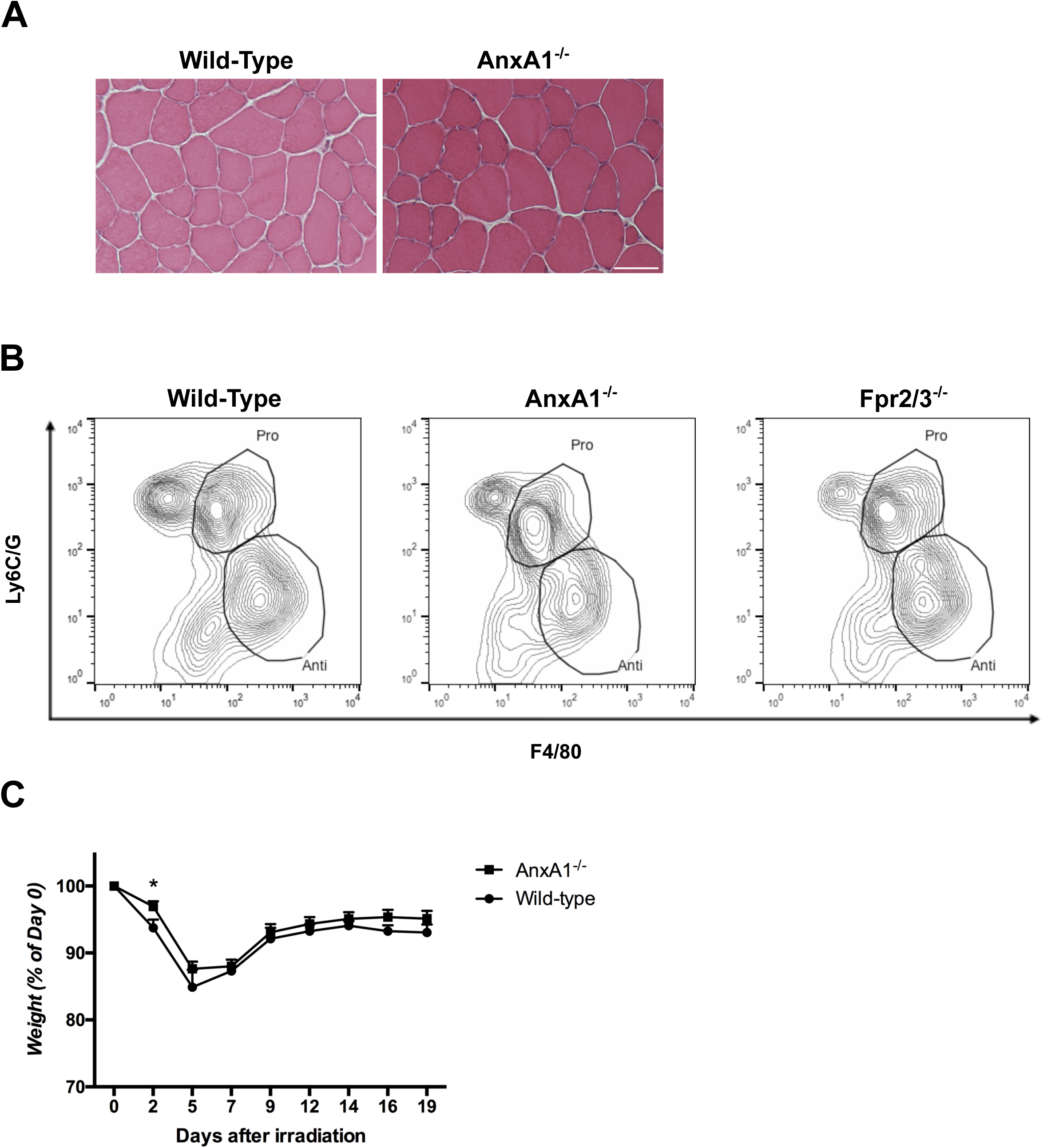
Related to Figure 1 and 2. (A) HE staining of non-injured Wild-Type and AnxA1^-/-^ *Tibialis Anterior* (TA) muscles. White bar = 50 μm. (B) Representative FACS plots of F4/80 and Ly6C/G markers in TA of Wild-Type, AnxA1^-/-^ and Fpr2/3^-/-^ mice 2 days post-cardiotoxin injury. (C) Weight follow-up, represented as percentage of Day 0, of mice after irradiation and bone marrow transplantation. Results are means ± SEM of at least six animals. *p<0.05 *versus* Day 0

**Suppl Fig 4.**
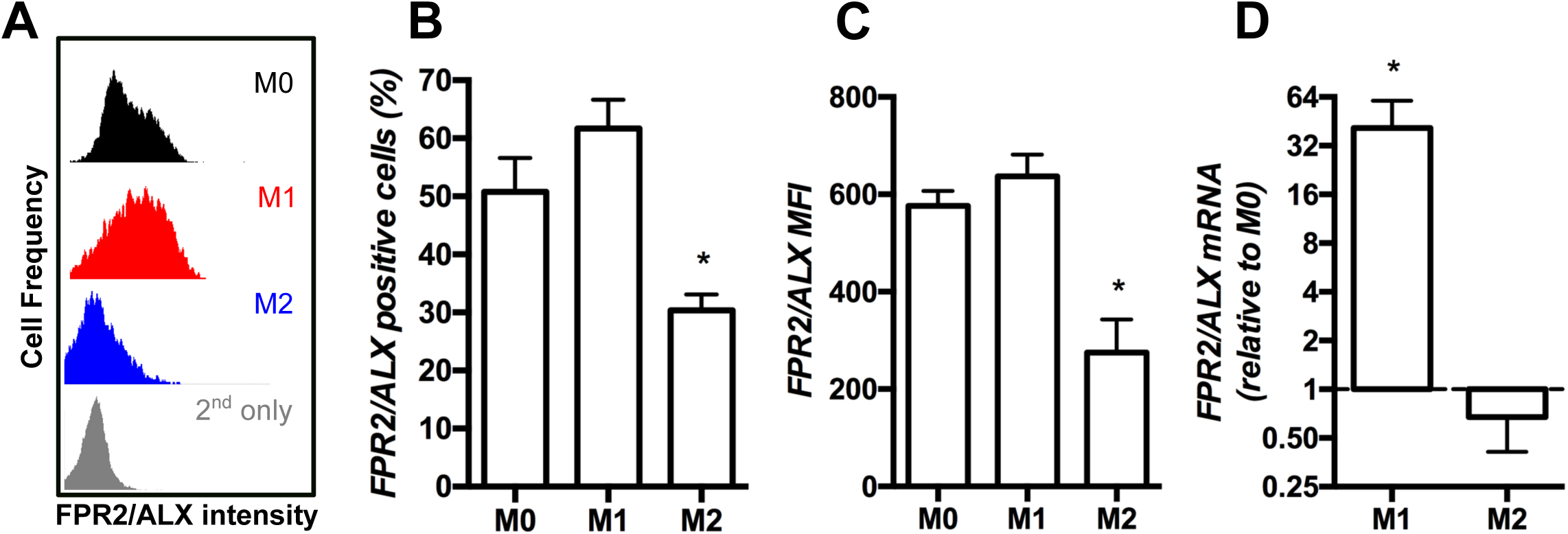
Related to Figure 3. FPR2/ALX expression varies following human macrophage polarization. Human primary macrophages were polarised into M1 or M2 macrophages with IFNg and IL-4, respectively (A-C) Flow cytometry analysis of FPR2/ALX expression. Shown are representative FACS plots (A), percentage of cells expressing FPR2/ALX (B) and FPR2/ALX MFI (C). (D) RT-qPCR analysis of FPR2/ALX mRNA level. Results are means +/- SEM of at least four independent experiments. *p < 0.05 *versus* M0 (non-activated).

## References

Amici, S.A., Dong, J., and Guerau-de-Arellano, M. (2017). Molecular Mechanisms Modulating the Phenotype of Macrophages and Microglia. Front Immunol 8, 1520.

Arnold, L., Henry, A., Poron, F., Baba-Amer, Y., van Rooijen, N., Plonquet, A., Gherardi, R.K., and Chazaud, B. (2007). Inflammatory monocytes recruited after skeletal muscle injury switch into antiinflammatory macrophages to support myogenesis. The Journal of experimental medicine 204, 1057–1069.

Bizzarro, V., Belvedere, R., Dal Piaz, F., Parente, L., and Petrella, A. (2012). Annexin A1 induces skeletal muscle cell migration acting through formyl peptide receptors. PLoS One 7, e48246.

Chan, K.L., Pillon, N.J., Sivaloganathan, D.M., Costford, S.R., Liu, Z., Theret, M., Chazaud, B., and Klip, A. (2015). Palmitoleate Reverses High Fat-induced Proinflammatory Macrophage Polarization via AMP-activated Protein Kinase (AMPK). The Journal of biological chemistry 290, 16979–16988.

d’Albis, A., Couteaux, R., Janmot, C., Roulet, A., and Mira, J.C. (1988). Regeneration after cardiotoxin injury of innervated and denervated slow and fast muscles of mammals. Myosin isoform analysis. Eur J Biochem 174, 103–110.

Dalli, J., Jones, C.P., Cavalcanti, D.M., Farsky, S.H., Perretti, M., and Rankin, S.M. (2012). Annexin A1 regulates neutrophil clearance by macrophages in the mouse bone marrow. FASEB J 26, 387–396.

Damazo, A.S., Yona, S., Flower, R.J., Perretti, M., and Oliani, S.M. (2006). Spatial and temporal profiles for anti-inflammatory gene expression in leukocytes during a resolving model of peritonitis. J Immunol 176, 4410–4418.

Davies, L.C., and Taylor, P.R. (2015). Tissue-resident macrophages: then and now. Immunology 144, 541–548.

Dufton, N., Hannon, R., Brancaleone, V., Dalli, J., Patel, H.B., Gray, M., D’Acquisto, F., Buckingham, J.C., Perretti, M., and Flower, R.J. (2010). Anti-inflammatory role of the murine formyl-peptide receptor 2: ligand-specific effects on leukocyte responses and experimental inflammation. J Immunol 184, 2611–2619.

Francis, J.W., Balazovich, K.J., Smolen, J.E., Margolis, D.I., and Boxer, L.A. (1992). Human neutrophil annexin I promotes granule aggregation and modulates Ca(2+)-dependent membrane fusion. J Clin Invest 90, 537–544.

Gobbetti, T., Coldewey, S.M., Chen, J., McArthur, S., le Faouder, P., Cenac, N., Flower, R.J., Thiemermann, C., and Perretti, M. (2014). Nonredundant protective properties of FPR2/ALX in polymicrobial murine sepsis. Proceedings of the National Academy of Sciences of the United States of America 111, 18685–18690.

Hannon, R., Croxtall, J.D., Getting, S.J., Roviezzo, F., Yona, S., Paul-Clark, M.J., Gavins, F.N., Perretti, M., Morris, J.F., Buckingham, J.C., et al. (2003). Aberrant inflammation and resistance to glucocorticoids in annexin 1-/- mouse. FASEB J 17, 253–255.

Jha, A.K., Huang, S.C., Sergushichev, A., Lampropoulou, V., Ivanova, Y., Loginicheva, E., Chmielewski, K., Stewart, K.M., Ashall, J., Everts, B., et al. (2015). Network integration of parallel metabolic and transcriptional data reveals metabolic modules that regulate macrophage polarization. Immunity 42, 419–430.

Jorgensen, S.B., Wojtaszewski, J.F., Viollet, B., Andreelli, F., Birk, J.B., Hellsten, Y., Schjerling, P., Vaulont, S., Neufer, P.D., Richter, E.A., et al. (2005). Effects of alpha-AMPK knockout on exercise-induced gene activation in mouse skeletal muscle. FASEB J 19, 1146–1148.

Kjobsted, R., Hingst, J.R., Fentz, J., Foretz, M., Sanz, M.N., Pehmoller, C., Shum, M., Marette, A., Mounier, R., Treebak, J.T., et al. (2018). AMPK in skeletal muscle function and metabolism. FASEB J 32, 1741–1777.

Lantier, L., Fentz, J., Mounier, R., Leclerc, J., Treebak, J.T., Pehmoller, C., Sanz, N., Sakakibara, I., Saint-Amand, E., Rimbaud, S., et al. (2014). AMPK controls exercise endurance, mitochondrial oxidative capacity, and skeletal muscle integrity. FASEB journal: official publication of the Federation of American Societies for Experimental Biology.

Lantier, L., Mounier, R., Leclerc, J., Pende, M., Foretz, M., and Viollet, B. (2010). Coordinated maintenance of muscle cell size control by AMP-activated protein kinase. FASEB journal: official publication of the Federation of American Societies for Experimental Biology 24, 3555–3561.

Leoni, G., and Nusrat, A. (2016). Annexin A1: shifting the balance towards resolution and repair. Biol Chem 397, 971–979.

Lim, L.H., and Pervaiz, S. (2007). Annexin 1: the new face of an old molecule. FASEB J 21, 968–975.

Locatelli, I., Sutti, S., Jindal, A., Vacchiano, M., Bozzola, C., Reutelingsperger, C., Kusters, D., Bena, S., Parola, M., Paternostro, C., et al. (2014). Endogenous annexin A1 is a novel protective determinant in nonalcoholic steatohepatitis in mice. Hepatology 60, 531–544.

Lucas, T., Waisman, A., Ranjan, R., Roes, J., Krieg, T., Muller, W., Roers, A., and Eming, S.A. (2010). Differential roles of macrophages in diverse phases of skin repair. J Immunol 184, 3964–3977.

Maderna, P., Yona, S., Perretti, M., and Godson, C. (2005). Modulation of phagocytosis of apoptotic neutrophils by supernatant from dexamethasone-treated macrophages and annexin-derived peptide Ac(2-26). J Immunol 174, 3727–3733.

Mantovani, A., Biswas, S.K., Galdiero, M.R., Sica, A., and Locati, M. (2013). Macrophage plasticity and polarization in tissue repair and remodelling. J Pathol 229, 176–185.

Martinez, F.O., and Gordon, S. (2014). The M1 and M2 paradigm of macrophage activation: time for reassessment. F1000Prime Rep 6, 13.

McArthur, S., Gobbetti, T., Kusters, D.H., Reutelingsperger, C.P., Flower, R.J., and Perretti, M. (2015). Definition of a Novel Pathway Centered on Lysophosphatidic Acid To Recruit Monocytes during the Resolution Phase of Tissue Inflammation. J Immunol 195, 1139–1151.

Moraes, L.A., Kar, S., Foo, S.L., Gu, T., Toh, Y.Q., Ampomah, P.B., Sachaphibulkij, K., Yap, G., Zharkova, O., Lukman, H.M., et al. (2017). Annexin-A1 enhances breast cancer growth and migration by promoting alternative macrophage polarization in the tumour microenvironment. Sci Rep 7, 17925.

Motwani, M.P., and Gilroy, D.W. (2015). Macrophage development and polarization in chronic inflammation. Semin Immunol 27, 257–266.

Mounier, R., Lantier, L., Leclerc, J., Sotiropoulos, A., Foretz, M., and Viollet, B. (2011). Antagonistic control of muscle cell size by AMPK and mTORC1. Cell cycle (Georgetown, Tex 10, 2640–2646.

Mounier, R., Lantier, L., Leclerc, J., Sotiropoulos, A., Pende, M., Daegelen, D., Sakamoto, K., Foretz, M., and Viollet, B. (2009). Important role for AMPKalpha1 in limiting skeletal muscle cell hypertrophy. FASEB journal: official publication of the Federation of American Societies for Experimental Biology 23, 2264–2273.

Mounier, R., Theret, M., Arnold, L., Cuvellier, S., Bultot, L., Goransson, O., Sanz, N., Ferry, A., Sakamoto, K., Foretz, M., et al. (2013). AMPKalpha1 Regulates Macrophage Skewing at the Time of Resolution of Inflammation during Skeletal Muscle Regeneration. Cell metabolism 18, 251–264.

Mounier, R., Theret, M., Lantier, L., Foretz, M., and Viollet, B. (2015). Expanding roles for AMPK in skeletal muscle plasticity. Trends Endocrinol Metab 26, 275–286.

Murray, P.J., Allen, J.E., Biswas, S.K., Fisher, E.A., Gilroy, D.W., Goerdt, S., Gordon, S., Hamilton, J.A., Ivashkiv, L.B., Lawrence, T., et al. (2014). Macrophage activation and polarization: nomenclature and experimental guidelines. Immunity 41, 14–20.

O’Neill, L.A., Kishton, R.J., and Rathmell, J. (2016). A guide to immunometabolism for immunologists. Nat Rev Immunol 16, 553–565.

Park, S.Y., Lee, S.W., Lee, S.Y., Hong, K.W., Bae, S.S., Kim, K., and Kim, C.D. (2017). SIRT1/Adenosine Monophosphate-Activated Protein Kinase alpha Signaling Enhances Macrophage Polarization to an Anti-inflammatory Phenotype in Rheumatoid Arthritis. Front Immunol 8, 1135.

Pearce, E.L., and Pearce, E.J. (2013). Metabolic pathways in immune cell activation and quiescence. Immunity 38, 633–643.

Perretti, M., Cooper, D., Dalli, J., and Norling, L.V. (2017). Immune resolution mechanisms in inflammatory arthritis. Nat Rev Rheumatol 13, 87–99.

Perretti, M., and D’Acquisto, F. (2009). Annexin A1 and glucocorticoids as effectors of the resolution of inflammation. Nat Rev Immunol 9, 62–70.

Perretti, M., Leroy, X., Bland, E.J., and Montero-Melendez, T. (2015). Resolution Pharmacology: Opportunities for Therapeutic Innovation in Inflammation. Trends Pharmacol Sci 36, 737–755.

Rhys, H.I., Dell’Accio, F., Pitzalis, C., Moore, A., Norling, L.V., and Perretti, M. (2018). Neutrophil Microvesicles from Healthy Control and Rheumatoid Arthritis Patients Prevent the Inflammatory Activation of Macrophages. EBioMedicine 29, 60–69.

Rodriguez-Prados, J.C., Traves, P.G., Cuenca, J., Rico, D., Aragones, J., Martin-Sanz, P., Cascante, M., and Bosca, L. (2010). Substrate fate in activated macrophages: a comparison between innate, classic, and alternative activation. J Immunol 185, 605–614.

Saclier, M., Cuvellier, S., Magnan, M., Mounier, R., and Chazaud, B. (2013a). Monocyte/macrophage interactions with myogenic precursor cells during skeletal muscle regeneration. The FEBS journal 280, 4118–4130.

Saclier, M., Yacoub-Youssef, H., Mackey, A.L., Arnold, L., Ardjoune, H., Magnan, M., Sailhan, F., Chelly, J., Pavlath, G.K., Mounier, R., et al. (2013b). Differentially activated macrophages orchestrate myogenic precursor cell fate during human skeletal muscle regeneration. Stem Cells 31, 384–396.

Scannell, M., Flanagan, M.B., deStefani, A., Wynne, K.J., Cagney, G., Godson, C., and Maderna, P. (2007). Annexin-1 and peptide derivatives are released by apoptotic cells and stimulate phagocytosis of apoptotic neutrophils by macrophages. J Immunol 178, 4595–4605.

Serhan, C.N. (2014). Pro-resolving lipid mediators are leads for resolution physiology. Nature 510, 92–101.

Serhan, C.N., and Savill, J. (2005). Resolution of inflammation: the beginning programs the end. Nat Immunol 6, 1191–1197.

Solito, E., Kamal, A., Russo-Marie, F., Buckingham, J.C., Marullo, S., and Perretti, M. (2003). A novel calcium-dependent proapoptotic effect of annexin 1 on human neutrophils. FASEB J 17, 1544–1546.

Tabas, I., and Glass, C.K. (2013). Anti-inflammatory therapy in chronic disease: challenges and opportunities. Science 339, 166–172.

Tannahill, G.M., Curtis, A.M., Adamik, J., Palsson-McDermott, E.M., McGettrick, A.F., Goel, G., Frezza, C., Bernard, N.J., Kelly, B., Foley, N.H., et al. (2013). Succinate is an inflammatory signal that induces IL-1beta through HIF-1alpha. Nature 496, 238–242.

Theret, M., Gsaier, L., Schaffer, B., Juban, G., Ben Larbi, S., Weiss-Gayet, M., Bultot, L., Collodet, C., Foretz, M., Desplanches, D., et al. (2017). AMPKalpha1-LDH pathway regulates muscle stem cell self-renewal by controlling metabolic homeostasis. Embo J 36, 1946–1962.

Troidl, C., Mollmann, H., Nef, H., Masseli, F., Voss, S., Szardien, S., Willmer, M., Rolf, A., Rixe, J., Troidl, K., et al. (2009). Classically and alternatively activated macrophages contribute to tissue remodelling after myocardial infarction. J Cell Mol Med 13, 3485–3496.

Van den Bossche, J., O’Neill, L.A., and Menon, D. (2017). Macrophage Immunometabolism: Where Are We (Going)? Trends Immunol 38, 395–406.

Varga, T., Mounier, R., Gogolak, P., Poliska, S., Chazaud, B., and Nagy, L. (2013). Tissue LyC6-macrophages are generated in the absence of circulating LyC6-monocytes and Nur77 in a model of muscle regeneration. J Immunol 191, 5695–5701.

Varga, T., Mounier, R., Horvath, A., Cuvellier, S., Dumont, F., Poliska, S., Ardjoune, H., Juban, G., Nagy, L., and Chazaud, B. (2016a). Highly Dynamic Transcriptional Signature of Distinct Macrophage Subsets during Sterile Inflammation, Resolution, and Tissue Repair. J Immunol 196, 4771–4782.

Varga, T., Mounier, R., Patsalos, A., Gogolak, P., Peloquin, M., Horvath, A., Pap, A., Daniel, B., Nagy, G., Pintye, E., et al. (2016b). Macrophage PPARgamma, a Lipid Activated Transcription Factor Controls the Growth Factor GDF3 and Skeletal Muscle Regeneration. Immunity 45, 1038–1051.

